# Study of In silico Anti-Inflammatory Potential of Tyramine-Fe complex in Brotowali Plants

**DOI:** 10.1101/2020.08.24.264473

**Authors:** T.W. Wimbuh, B.S. Sutiman, Sri Widyarti, H. S Djoko

## Abstract

Tyramine-Fe complex is a bioinorganic complex that is formed in brotowali. The complex is thought to play a role in the anti-inflammatory activity of brotowali. The mechanism of anti-inflammatory drugs such as aspirin and ibuprofen is to inhibit the reaction of prostaglandin formation from COX2 and arachidonic acid. In this study, a docking simulation was performed between COX2 with single tyramine, tyramine-Fe complex ibuprofen, and aspirin. The results showed that the COX2-single tyramine bond overlapped with COX2-aspirin, while COX2-tyramine-Fe complex bond overlapped with COX2-ibuprofen. The energy required to form COX2-tyramine-Fe complex bond was smaller than that of single tyramine, aspirin, and ibuprofen. The number of bonds of COX2-tyramine-Fe complex bond is higher than that of single tyramine. These results proved that COX2-tyramine-Fe complex bond had anti-inflammatory activity and greater activity than single tyramine.

## Introduction

Inflammation is part of the complex biological response of body tissues to harmful stimuli, such as pathogens, damaged cells, or irritants (Ferrero-Miliani et al. 2007). The function of inflammation is to eliminate the initial cause of cell injury, clear out necrotic cells and tissues damaged from the original insult and the inflammatory process, and initiate tissue repair. However, progressive inflammation can cause certain unwanted diseases, such as fever, periodontitis, atherosclerosis, rheumatoid arthritis, and even cancer (Chen et al. 2018).

Aspirin and ibuprofen are widely used anti-inflammatory drugs. Both are included in the class of Nonsteroidal Anti-inflammatory Drugs (NSAIDs). NSAIDs’ mechanism is to block the reaction between arachidonic acid and the COX enzyme so that prostaglandins are not formed, which are mediators of inflammation (Ricciotti and Fitzgerald 2011). However, NSAIDs have several side effects, including gastrointestinal damage, renal syndrome, and respiratory effect (Wongrakpanich et al. 2018). So we need anti-inflammatory alternatives that have minor side effects. Alkanoids contained in herbal plants have anti-inflammatory activities (Souto et al. 2011).

Brotowali is an herb that contains flavonoids, terpenoids, alkaloids, lignans, nucleosides, and sterols. The content of various compounds in brotowali is closely related to its pharmacological activity (Ahmad, Jantan, and Bukhari 2016). Brotowali is widely used by traditional communities in various countries, including in Indonesia (Pathak, Jain, and Sharma 1995; Dweck and Cavin J. P. 2006), Malaysia (Rahman et al. 1999), Thailand (Rungruang and Boonmars 2009), (Quisumbing 1951) and several other countries that have tropical and sub-tropical climates. Several studies have proven that Brotowali has anti-inflammatory activity (Kamarazaman et al. 2012; Hipol, Cariaga, and Hipol 2012) so that it can be used as an alternative to NSAIDs. One of the active compounds brotowali from the alkaloids group is tyramine. Apart from brotowali, tyramine is also found in various Cactaceae and in the Leguminosae (Smith 1977). Tyramine and Fe in brotowali are predicted forming a bioinorganic complex.

Several studies have shown that the transition metals present in plants can form bioinorganic complexes. These studies include the results of research from Sukmaningsih et al. (2018), which proved the flavonoid complex with Fe in juwet plants. Capo et al. (2017) prove that the Oleuropein complex with copper is also found in the leaves, stems, and processed products of Olives (Olea europea). Transition metals in plants come from transition metals that are absorbed by plant roots from the soil.

COX2 is an enzyme that catalyzes the reaction of prostaglandin formation from arachidonic acid during infection (Zarghi and Arfaei 2011). In this research, a docking simulation will be performed between single tyramine and tyramine-Fe complexes with COX2. The results of this analysis are expected to provide an overview of the anti-inflammatory potential of the tyramine-Fe complex contained in Brotowali.

## Methods

The structure of the tyramine-Fe complex was drawn with the ChemDraw Pro 8.0 application. Docking was done with the AutoDock Vina 1.1 program in the Pyrx program. Docking simulations were performed between COX2-tyramine, COX2-tyramine-Fe complexes, COX2-aspirin, and COX2-ibuprofen. The 3D structure of the COX2 protein (PDB ID: 5f19) was downloaded from the RCSB database (www.rcsb.org). The structures of tyramine (PubChem ID: 5610), aspirin (PubChem ID: 2244), and ibuprofen (PubChem ID: 3672) were downloaded from the PubChem database (www.pubchem.ncbi.nlm.nih.gov). Visualization and interpretation of docking results using the Pymol v.2.3.2.1 and Discovery Studio 2019 programs.

## Results and Discussion

Fe is a transition metal that can bind several ligands, while tyramine is an alkaloid that has one –OH group (Figure 1). The metal Fe and the –OH group on the tyramine can bind to form the tyramine-Fe complex. One possibility of the complex structure formed is that one Fe binds to 2 tyramine molecules (Figure 2).

**Figure 1.**
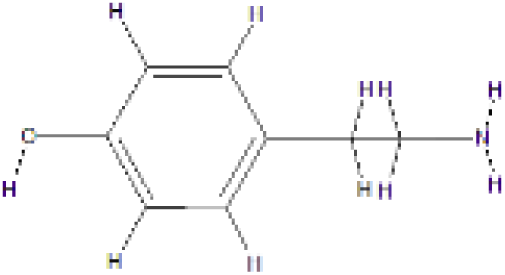
Single tyramine structure.

**Figure 2.**
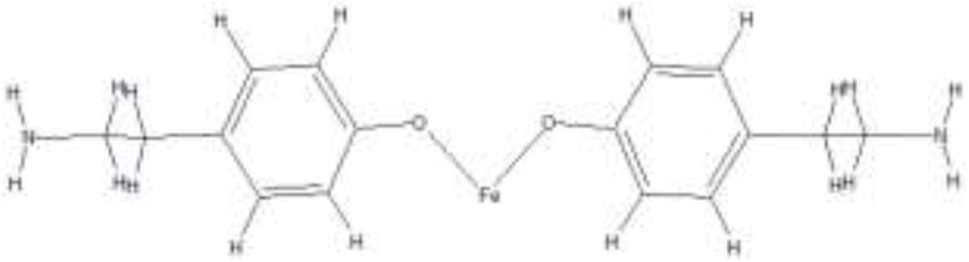
Structure of tyramine-Fe complex.

The results of COX2 docking with tyramine showed that the COX2-tyramine bond overlaps with the COX2-Aspirin bond site (Figure 3). COX2 bonds with tyramine to form hydrogen bonds on amino acid residues GLN203, ALA199, TYR385, and hydrophobic bonds on amino acid residues GLN203, ALA202. The COX2 bonds with tyramine form hydrogen bonds in the amino acid residues TRP387, HIS386, and form hydrophobic bonds at the amino acid residues HIS388, ALA202. The energy required to form a COX2-tyramine bond is -5.6 kcal/mol, while the energy required to form a COX2-aspirin bond is -6.4 kcal/mol (Table.1).

**Tabel 1.** Binding Affinity.

| Interactions | Binding Affinity (kcal/mol) |
| --- | --- |
| COX2-tyramine | -5.6 |
| COX2-aspirin | -6.4 |
| COX2- tyramine-Fe complex | -7.4 |
| COX2- ibuprofen | -6.6 |

**Figure 3.**
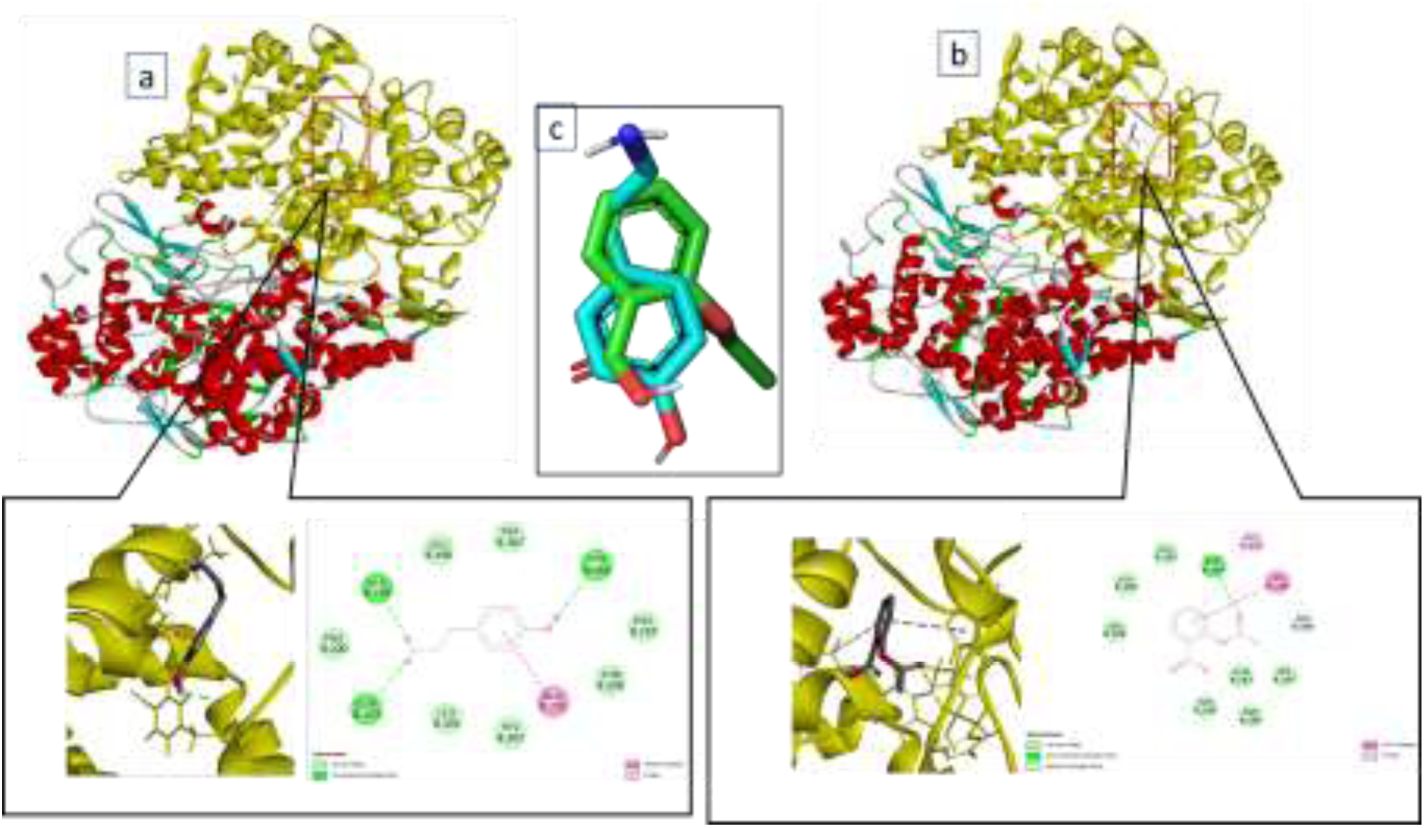
Comparison of docking results between COX2-Tyramine and COX2-Aspirin; (a) bond position between COX2-Tyramine, (b) bond position between and COX2-Aspirin, (C) Overlapping position of Tyramine (blue) and Aspirin (green) on COX2.

The results of COX2 docking with the tyramine-Fe complex showed that the COX2-tyramine-Fe complex overlapped with the COX2-ibuprofen bond sites (Figure 4). The COX2 bonds with the tyramine-Fe complex form hydrogen bonds on the amino acid residues of CYS41, GLN461, and hydrophobic bonds on the amino acid residues of CYS36, CYS47, PRO153, PRO156, LEU152. COX2 bonds with ibuprofen to form hydrogen bonds in the amino acid residue ASN43 and hydrophobic bonds on amino acid residues LEU152, LYS468, ARG469, PRO153, CYS36, PRO40, CYS47, LEU152, PRO153, HIS39. The energy required to form a COX2-tyramine-Fe bond is -7.4 kcal/mol, while the energy required to form a COX2-ibuprofen bond is - 6.6 kcal/mol.

**Figure 4.**
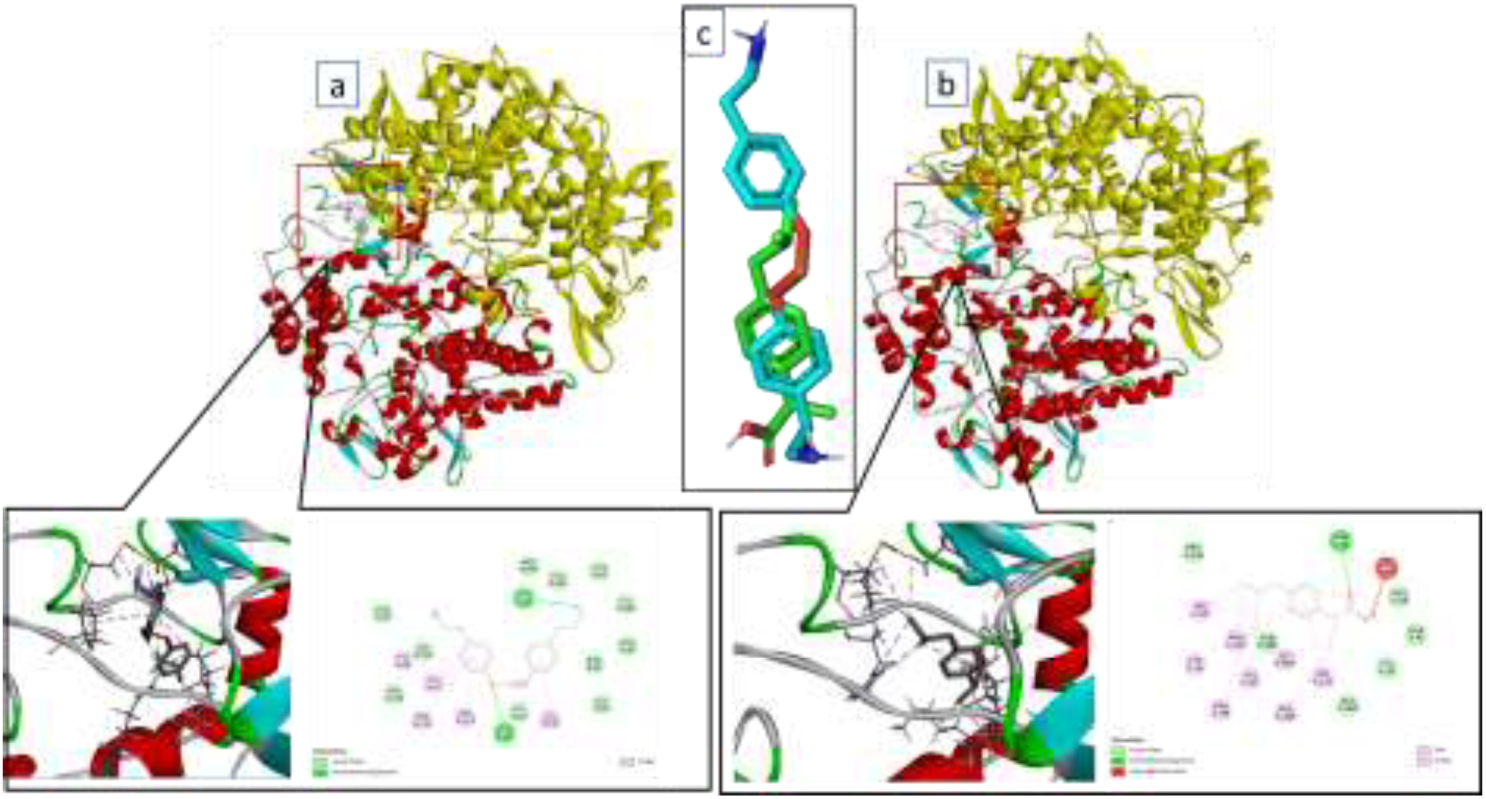
Comparison of docking results between COX2-Tyramine complex with Fe and COX2-ibuprofen; (a) bond position between COX2-Tyramine complex with Fe, (b) bond position between COX2-ibuprofen, (C) Overlapping position of Tyramine complex with Fe (blue) and ibuprofen (green) on COX2

Aspirin and ibuprofen are NSAIDs. Overlapping of COX2-tyramine and COX2-tyramine-Fe complex proves that tyramine and tyramine-Fe complexes can act as an anti-inflammatory as in aspirin and ibuprofen. However, the energy required by the tyramine-Fe complex is smaller than the single tyramine. The lower the energy required, the easier the bonds are to form so that the tyramine-Fe complex binds more easily with COX2 compared to single tyramine.

When interacted with COX2, single tyramine required more energy than aspirin. However, when it has formed a tyramine-Fe complex, the energy required is smaller than ibuprofen, aspirin, and single tyramine. The number of bonds formed between COX2 and the tyramine-Fe complex was greater than that formed between COX2 and single tyramine. The tyramine-Fe complex forms two hydrogen bonds and five hydrophobic bonds. In contrast, single tyramine forms three hydrogen bonds and two hydrophobic bonds.The lower energy required for bonding and the more bonds formed between COX2-tyramine-Fe complex indicates that the tyramine-Fe complex has better anti-inflammatory activity than the single tyramine.

Anti-inflammatory drugs such as aspirin and ibuprofen only have a single role, which is to prevent prostaglandin formation reactions so that there is no inflammatory reaction. Excessive consumption of anti-inflammatory drugs also causes stomach ulcers.

Brotowali has many compounds. Apart from the tyramine-Fe complex, which has an anti-inflammatory role, other compounds act as antioxidants, immunomodulators, anticholinesterases, and antimalarials. Therefore, consumption of herbal herbs, including brotowali, is highly recommended for people with infectious diseases and to maintain fitness, especially herbal medicine has been trusted and has been proven to have very little side effects.

## Conclusion

The potential of anti-inflammatory activity of tyramine-Fe complex in brotowali is better than single tyramine, aspirin and ibuprofen provide new alternative options in the treatment, cure, and prevention of infectious diseases.

## Suggestion

The structure of the tyramine-Fe complex used for docking simulations should be processed first in the Avogadro program to obtain the optimal structure. In this study, tyramine-Fe complex structure, which was drawn using the ChemDraw Pro 8.0 program, was immediately used as a ligand.

